# Reduced metabolic activity in established *Gardnerella* spp. biofilms contributes to protection from the bactericidal effects of metronidazole

**DOI:** 10.1101/2021.09.06.459156

**Authors:** Salahuddin Khan, Janet E. Hill

**Affiliations:** Department of Veterinary Microbiology, University of Saskatchewan, Saskatoon, Saskatchewan, Canada, S7N 5B4

## Abstract

*Gardnerella* spp. can form biofilm and it has been suggested that failure of antibiotic treatment of bacterial vaginosis and recurrent vaginosis are linked to this property, however no specific mechanisms have been identified. Here, we performed broth microdilution assays to measure the minimum inhibitory concentration (MIC) of metronidazole for thirty-five *Gardnerella* isolates in one medium in which *Gardnerella* spp. grow primarily as biofilm and another in which growth is primarily planktonic. The MIC of isolates in the two conditions were highly correlated (R^2^= 0.69, p <0.001). No significant reduction in viable cell count of 7/9 isolates was observed when established biofilms were exposed to metronidazole at levels double the MIC. Subsequent measurements of metabolic activity in established biofilms formed by a larger collection of 31 isolates showed reduced metabolic activity following treatment with 128 μg/ml of metronidazole relative to untreated controls in 27/31 cases. The amount of biofilm produced by *Gardnerella* isolates was not enhanced by metronidazole exposure, even at sub-MIC levels. Our results suggest that reduced metabolic activity in *Gardnerella* spp. growing in established biofilms may be a mechanism of protection from the bactericidal effects of metronidazole.

## Introduction

*Gardnerella* spp. are considered a hallmark of bacterial vaginosis: a vaginal dysbiosis defined by the shift from a *Lactobacillus* spp. dominated microbiome to a more diverse microbiome, comprising many aerobic and anaerobic bacteria, including *Gardnerella* spp.. The genus *Gardnerella* includes at least four cpn60-defined subgroups corresponding to four whole-genome sequence defined clades, which were more recently amended to thirteen genome species (Ahmed *et al*. 2012; Paramel Jayaprakash, Schellenberg and Hill 2012; Vaneechoutte *et al*. 2019). One major diagnostic feature of bacterial vaginosis is the presence of clue cells – epithelial cells coated with multispecies biofilm (Swidsinski *et al*. 2005, 2013). It has been observed that *Gardnerella* accounts for a significant proportion of this multispecies biofilm *in vivo* (Hardy *et al*. 2017), and several studies have shown that *Gardnerella* spp. can also form biofilm *in vitro* (Harwich *et al*. 2010; Patterson *et al*. 2010; Machado *et al*. 2015; Khan, Voordouw and Hill 2019). Since the recognition of multiple *Gardnerella* spp., it has also been demonstrated *in vitro* that *Gardnerella* can form multispecies biofilm (Khan, Voordouw and Hill 2019). Biofilm formation is often considered a stress response that protects bacterial cells from environmental stresses such as antimicrobial substances, immune cells, and predators (Donlan and Costerton 2002; Jefferson 2004; Lewis 2008; Raghupathi *et al*. 2018; Vidakovic *et al*. 2018). Alternatively, biofilm may be a natural and preferred mode of growth for some bacteria in particular environments (Jefferson 2004).

Growth in biofilm form can provide protection from antibiotics in a variety of ways including enhanced production of extracellular matrix, reduction of metabolic activity, generation of environmental heterogeneity, and induction of phenotypic diversity (Mah and O’Toole 2001; Lebeaux, Ghigo and Beloin 2014). Slow diffusion of antimicrobials within a biofilm may allow the bacteria living within to produce more extracellular matrix to prevent penetration of antibiotics (Mah 2012). In fact, for some well characterized bacterial species, sublethal concentration of antibiotics have been demonstrated to lead to increased production of biofilm (Hoffman *et al*. 2005; Andersson and Hughes 2014; Oliveira *et al*. 2015a; Yu, Hallinen and Wood 2018).

Metronidazole, a bactericidal drug that causes DNA breakage by formation of reactive oxygen species (Sigeti, Guiney and Davis 1983), is widely prescribed for the treatment of bacterial vaginosis (Machado *et al*. 2015) and there is increasing concern about treatment failure (Swidsinski *et al*. 2008; Verwijs *et al*. 2020). Resistance to metronidazole can occur through reduced uptake of the prodrug, increased export and nitroimidazole reductase activity (Smith 2018; Alauzet, Lozniewski and Marchandin 2019). The ability of *Gardnerella* spp. to form biofilm is often suggested to be correlated with metronidazole treatment failure and recurrent BV (Swidsinski *et al*. 2008; Machado *et al*. 2015; Muzny and Schwebke 2015; Verwijs *et al*. 2020). In one recent study of a small number of *G. vaginalis* isolates the authors reported that the MIC of metronidazole was higher for biofilm forming isolates but whether this is true across all species of *Gardnerella* is not known, and no specific mechanism for the observed difference in MIC was suggested (Li et al. 2020). The objectives of our current study, therefore, were to determine if biofilm formation by *Gardnerella* spp. provides protection against metronidazole, if sub-inhibitory concentrations of metronidazole stimulate biofilm production, and to identify possible mechanisms of protection against metronidazole treatment of established *Gardnerella* biofilm.

## Methods

### Bacterial Isolates

Thirty-five *Gardnerella* isolates representing the four subgroups defined by cpn60 barcode sequence (n*=* 9, subgroup A (including *G. swidsinskii* and *G. leopoldii*); *n=* 10, subgroup B (*G. piotii* and genome sp. 3); *n* = 9 subgroup C (*G. vaginalis*); and *n=* 7, subgroup D (genome sp. 8, 9 and 10) were revived from freezer stocks on Columbia agar plates supplemented with 5% (v/v) sheep blood by incubating them anaerobically for 48 h at 37 ºC (BD GasPak EZ Anaerobe Gas Generating Pouch System, NJ, USA). To prepare inoculum for the broth microdilution assay, approximately ten well-isolated colonies from each blood agar plate were transferred to 5 ml brain heart infusion (BHI) medium supplemented with 0.25% (w/v) maltose and 10% (v/v) heat inactivated horse serum and incubated anaerobically for 18 h at 37 ºC.

### Broth microdilution assay

A stock solution, of metronidazole (102.4 mg/ml, M3761-25G, Sigma-Aldrich, ON, Canada) in DMSO was prepared and stored at -20 ºC. Immediately before each experiment, the stock solution was diluted 1:100 in DMSO to make a 1024 μg/ml solution. A broth microdilution assay was performed to determine the minimum inhibitory concentration (MIC) of metronidazole (Wiegand, Hilpert and Hancock 2008). Briefly, 100 μl of media were aliquoted into each well of a flat bottom 96 well plate (Corning Costar, NY, USA) using a multichannel pipettor. To make two-fold serial dilutions, 100 μl of 1024 μg/ml metronidazole was added to each well of the first column of a 96-well plate. After mixing by pipetting up and down 4-6 times, 100 μl was transferred to the second column, and the process was repeated to column 10. After pipetting up and down, 100 μl from column 10, instead of transferring to column 11, was discarded. Column 11 was used as growth control (no antibiotic). Column 12 was used negative control (sterile media). A freshly grown broth culture was adjusted to an OD_595_ of 0.5, corresponding to 10^6^-10^7^ cfu/ml, and 5 μl of adjusted broth culture was added to each well of the 96 well plate, up to column 11. The plates were then incubated anaerobically at 37 °C for 72 h. The process was repeated for all 35 isolates, and each isolate was tested in two growth media: BHI + 0.25% maltose (v/v) and BHI + 0.25% maltose (v/v) + 10% heat inactivated horse serum.

### Quantification of planktonic and biofilm growth

Following incubation of cultures in 96-well plates, the supernatant portion (planktonic growth) from each well was transferred to a fresh flat-bottom 96 well plate scanned at 595 nm using a microplate reader (VarioSkan LUX Multimode plate reader). Planktonic growth was reported as the OD_595_. Quantification of the biofilm remaining in each well after removal of the supernatant portion was performed using a CV assay as described previously (Khan, Voordouw and Hill 2019). Briefly, the 96-well plates were thoroughly washed twice with water. The wells were then stained with 1% (w/v) crystal violet for 20 minutes. Then the plates were washed twice with water and were dried before the addition of 33% (v/v) glacial acetic for biofilm solubilization. The plates were then read at 595 nm to quantify biofilm, which was reported as OD_595_ of crystal violet stain.

### Metabolic activity and viability measurement

Bacterial colonies harvested from blood agar plates were transferred to BHI + 0.25% maltose + 10% heat-inactivated horse blood serum and were incubated anaerobically for 18h. Inoculum was prepared using 18 h old culture by adjusting the OD to 0.5 (corresponding to 10^6^-10^7^ cfu/ml) in BHI + 0.25% maltose and 100 μl was pipetted into each well of duplicate 96-well plates (one plate for viability assay and one for metabolic activity measurement). One column of each plate was maintained containing sterile media only as negative control. After 48h of anaerobic incubation at 37 °C, each plate was divided into two sections: rows A through D as control and E through H as treatment. Overall OD_595_ was recorded, and the planktonic portion was transferred to a new plate to measure planktonic OD_595_ only. Fresh BHI + 0.25% maltose was pipetted into the control wells. For the treatment wells, fresh BHI + 0.25% maltose was supplemented with metronidazole at a final concentration of 128 μg/ml. Media supplemented with metronidazole were pipetted into treatment wells (Rows E-H) and the 96-well plates were incubated anaerobically at 37 °C for another 24h. Following 24h incubation, overall OD_595_ was measured. The planktonic fraction (supernatant) was transferred to a new plate and OD_595_ was recorded.

To determine viability of the cells growing in the biofilm mode post-treatment, biofilm formed at the bottom of wells were scraped into 100 μl of PBS added to each well, pipetted up and down to completely resuspend the cells and then 10 μl was transferred into new 96-well plate containing 90 μl of PBS using a multichannel pipettor. A serial dilution was made in the fresh 96-well plate and 10^−3^ through 10^−6^ dilutions were spotted (2.5 μl each spot) onto blood agar plates. The blood agar plates were incubated for 48 h and the colonies were counted on each spot. Each experiment contained four technical replicates.

To measure metabolic activity, 100 μl of PBS (pH 7) was added to each well of the original plate after removal of the planktonic growth. CellTitre Blue® reagent (20 μl) (Promega G8080) was pipetted into both plates: biofilm growth plate and planktonic growth plate, avoiding direct exposure of light. The incubation period for planktonic and biofilm fractions varied: visual colour change was observed in planktonic fractions containing plates sooner (at 20 min) than the biofilm cells containing plates. Plates were monitored every 20 min for up to 2 hours of incubation at 37 °C. Fluorescence for the biofilm forming cells was recorded after 2 h. Fluorescence was measured using VarioSkan LUX Multimode plate reader at 560 nm (excitation) and 590 nm (emission). Metabolic activity was measured by subtracting RFU values of the sterile control wells from the RFU values of the test wells.

### Statistical analysis

To test if biofilm growth and planktonic growth were significantly different in the two different media (with or without serum), a Mann-Whitney test was performed. To determine the relationship between the MIC values for *Gardnerella* isolates in the two different culture conditions, a Pearson-coefficient test was performed. To determine if the cell counts were significantly different between controls and treatments, a Mann-Whitney U test and Holm-Šídák multiple comparisons were performed. To test if metabolic activity was significantly different between treated and control biofilms, a multiple unpaired t-test and Holm-Šídák multiple comparisons were performed. All statistical analyses were performed using GraphPad Prism (v.9.2.0).

## Results

### Impact of serum on biofilm formation

We had previously determined that all *Gardnerella* spp. can form biofilm in culture to some extent (Paramel Jayaprakash, Schellenberg and Hill 2012; Khan, Voordouw and Hill 2019) and subsequently observed that the inclusion of serum in growth medium was associated with reduced biofilm formation. This offered an opportunity to control the growth mode of isolates, facilitating experiments investigating directly the relationship of growth mode and effects of metronidazole. To confirm these unpublished observations and quantify the effect of the presence of serum on biofilm formation, all isolates were grown in BHI + 0.25% (w/v) maltose with or without addition of 10% (v/v) heat inactivated horse serum. Thirty-five isolates were grown in four technical replicates. Overall, planktonic growth was significantly higher in media containing serum compared to serum-free media while biofilm growth was significantly higher in serum-free media than in media with serum (p <0.0001, Mann-Whitney U test) (Fig 1). On an individual basis, most isolates (74.28%, n= 26/35) had a significantly higher amount of biofilm formation in the media without serum (BHI+0.25% maltose (w/v)) than in the media with serum (BHI+0.25% maltose (w/v) + 10% horse serum), while planktonic growth was significantly higher in media with serum for majority of the isolates (57%, n= 20/35) (Fig S1).

**Fig 1:**
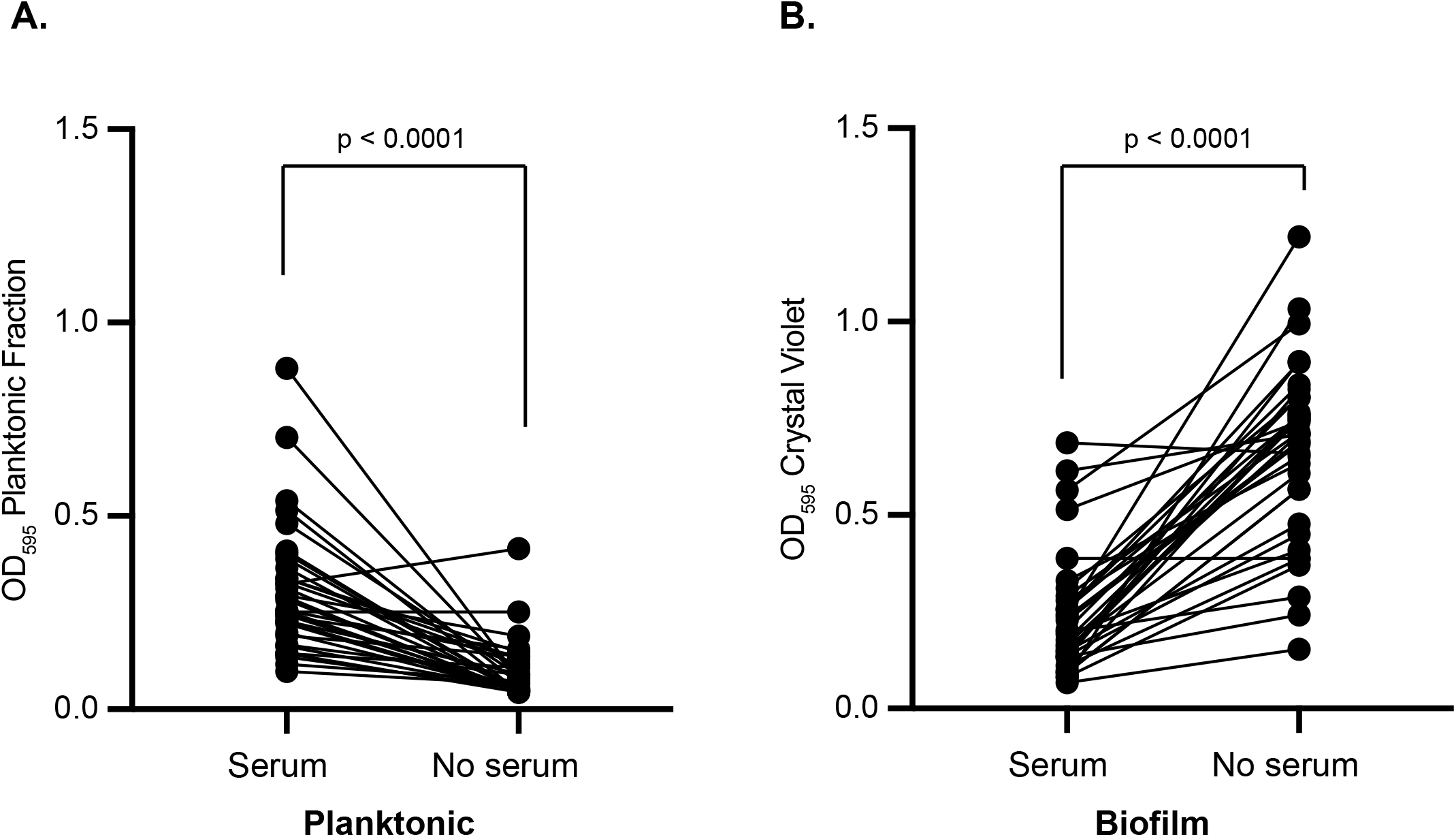
Growth mode is affected by serum. (a) Planktonic growth in media with serum (BHI+ 0.25% maltose (w/v) + 10% (v/v) heat inactivated horse serum) and without serum (BHI+ 0.25% (w/v) maltose) measured by absorbance at 595 nm. (B) Biofilm formation in media with serum (BHI+ 0.25% (w/v) maltose + 10% (v/v) heat inactivated horse serum) and without serum (BHI+ 0.25% (w/v) maltose) measured by crystal violet stain. Each data point is the average of four technical replicates of each of 35 isolates. Results for each isolate in the two conditions are connected by lines. Mann-Whitney test was performed to test significance and p values are indicated.

### Impact of growth mode on susceptibility to metronidazole

Since serum significantly reduces biofilm formation and encourages planktonic growth, we used presence or absence of serum to control mode of growth of the tested isolates in subsequent broth microdilution assays. If growth in biofilm reduces susceptibility to metronidazole, we would expect to see higher MIC values for isolates grown in serum-free media than in media with serum. Thirty-five *Gardnerella* isolates were grown in metronidazole concentrations ranging from 1 μg/ml to 512 μg/ml (two-fold serial dilution) in BHI + 0.25% (w/v) maltose with or without 10% (v/v) heat inactivated horse serum. The MIC for most of the tested isolates (19/35, 54.28%) was 64 μg/ml (range 4-128 μg/ml). Overall, MIC values determined in the two media were highly correlated (Pearson correlation co-efficient, R^2^= 0.69, p <0.001) and 27/35 isolates had the same MIC in both media. Of the 35 isolates tested, the observed MIC differed between the two conditions for eight (8/35, 23%) isolates: NR010 of subgroup A; GH007, GH019 & GH022 of subgroup B; and N165, GH015, GH021, and VN001 of subgroup C. Except N165, the seven other isolates (GH007, GH015, GH019, GH021, GH022, NR010, and VN001) had a lower MIC value in serum free media than in media with serum. There was no apparent relationship between subgroup and MIC (Table 1).

**Table 1.**
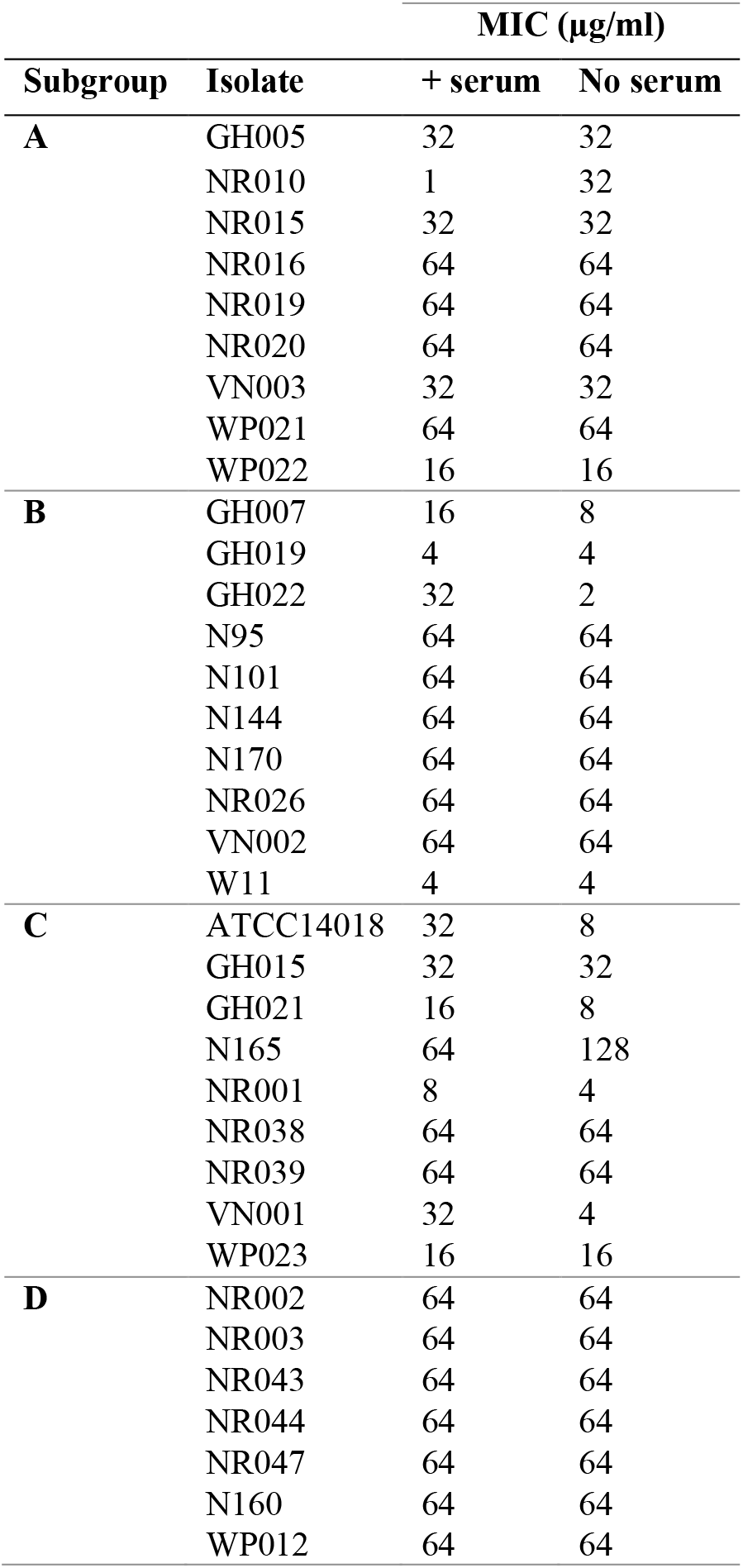
MIC of thirty-five *Gardnerella* isolates in BHI+ 0.25% (w/v) maltose with or without 10% (v/v) heat inactivated horse serum.

### Viability of treated biofilm vs. control biofilm

In broth microdilution assay, a suspension of cells are exposed to antibiotic simultaneously with inoculation of assay plates, which means that the bacteria are exposed to the antibiotic before they have the opportunity to establish biofilm. To investigate if established (preformed) biofilms formed by *Gardnerella* provide protection against subsequent metronidazole treatment, preformed biofilms (48 h old) of a subset of nine *Gardnerella* isolates were treated for 24 h with 128 μg/ml metronidazole, which is double the highest recorded MIC value for any of the tested isolates. Total viable counts of isolates treated with metronidazole and untreated controls were compared. Viable cells were recovered from all controls and all treated biofilms except VN003 and ATCC 14018, which had no viable cells after treatment. In cases where viable cells were recovered, there were no significant differences in cfu/ml between treated and untreated (Fig 2, Mann-Whitney test, Holm-Šídák multiple comparisons).

**Fig 2:**
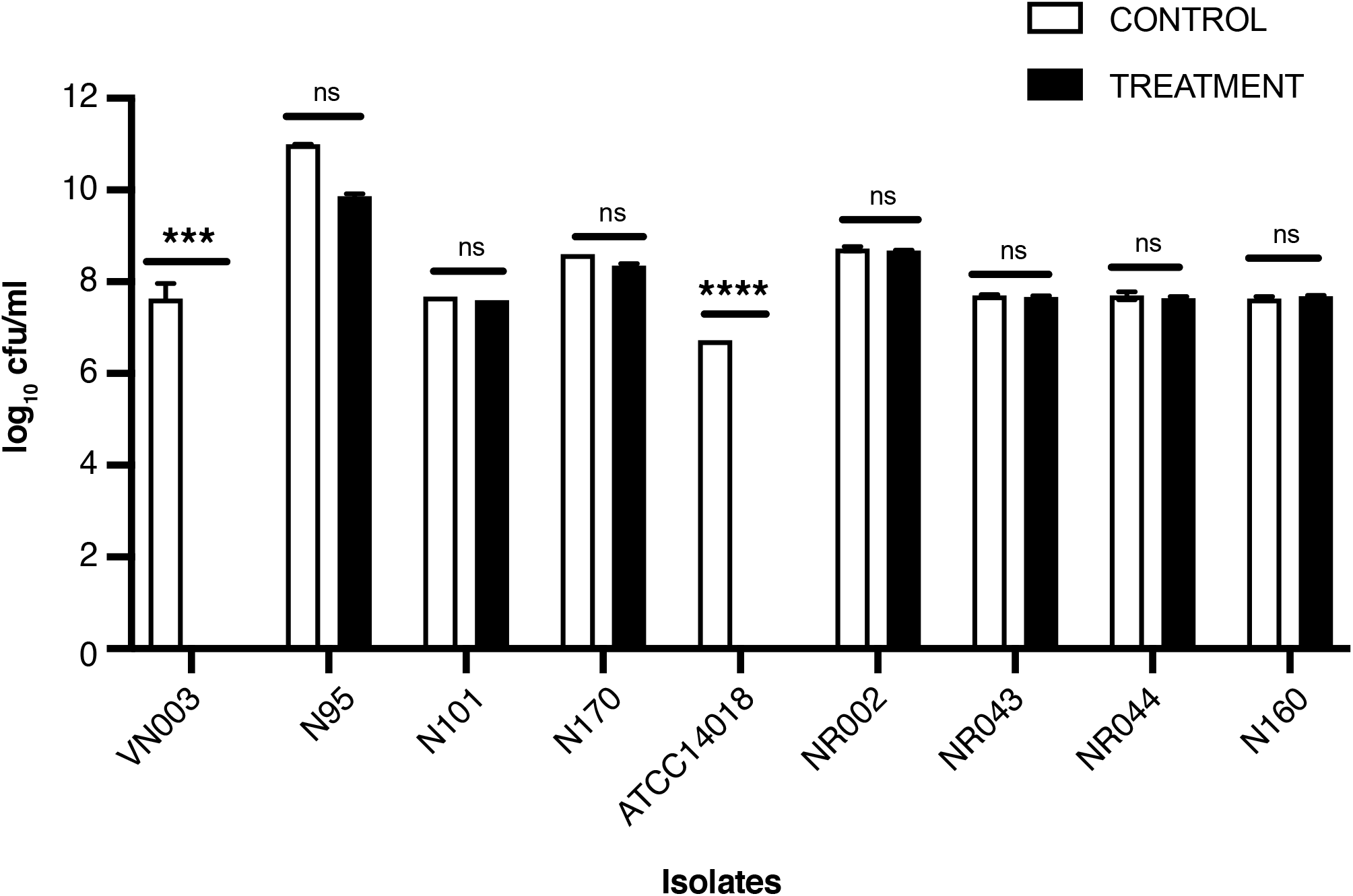
*Gardnerella* isolates in established biofilms can survive metronidazole treatment. 48 h old established biofilms of nine *Gardnerella* isolates were either treated with 128 μg/ml of metronidazole or not treated (controls). Biofilm cells were scraped, resuspended and total viable counts were performed on blood agar plates. Error bars show standard deviation of four technical replicates. Mann-Whitney U test was performed to test statistical significance (**** = p<0.0001, *** = p <0.001, ns = not significant).

### Metabolic activity of treated biofilm vs. control biofilm

To determine if treated and untreated biofilms differ in metabolic activity, a resazurin assay was performed. If cells are metabolically active, resazurin is reduced to resosurfin – a highly fluorescence substance. We measured fluorescence (560/590nm) for thirty-one *Gardnerella* isolates growing in biofilm mode, which were untreated or treated with 128 μg/ml metronidazole for 24 h. Metabolic activity, measured as RFU values, was appreciably higher in untreated biofilms: in 87% (27/31) of isolates metabolic activity was lower in the metronidazole treated biofilm than in the corresponding untreated control (Fig 3). Of the 27 isolates which showed reduced metabolic activity in preformed biofilms treated with metronidazole, the reduction was significant in seventeen isolates (Fig 3, unpaired t-test, Holm-Šídák multiple comparisons). Although three isolates (NR021, N95, NR026) showed higher metabolic activity in treatment than in controls, these differences were not significant (Fig 3).

**Fig 3:**
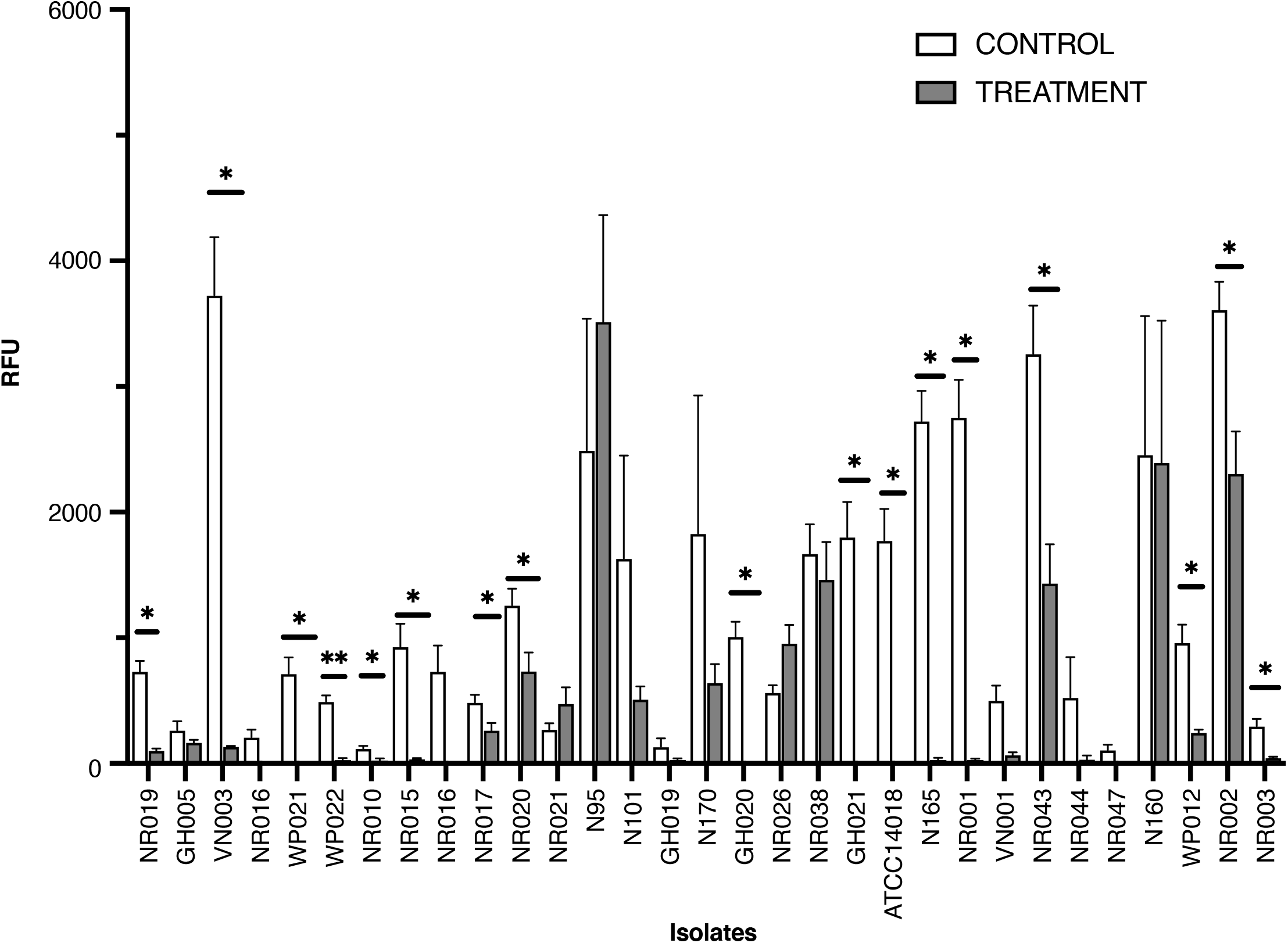
Metabolic activity of *Gardnerella* isolates growing in biofilms is reduced after metronidazole treatment. Thirty-three *Gardnerella* isolates were grown in BHI + 0.25% maltose for 48 h to form biofilms. Biofilms were treated with 128 μg/ml of metronidazole for 24 h, with four replicates per isolate. Metabolic activity was measured as baseline-subtracted relative fluorescence units (RFU) at 560/590 nm. An unpaired t-test and Holm-Šídák multiple comparisons were performed to test if the differences were statistically significant (* = p <0.005, ** = p <0.05).

### Impact of metronidazole concentration on biofilm formation

To investigate if metronidazole at any concentration could stimulate biofilm formation by *Gardnerella*, we compared the amount of biofilm growth of each isolate at each concentration of metronidazole in serum-free media. No enhancement of biofilm formation was observed in most of the tested isolates in the presence of metronidazole compared to biofilm formation in the absence of metronidazole, except for NR026 (Subgroup B), VN001 (Subgroup C), and NR043 (Subgroup D) (Fig 4).

**Fig 4:**
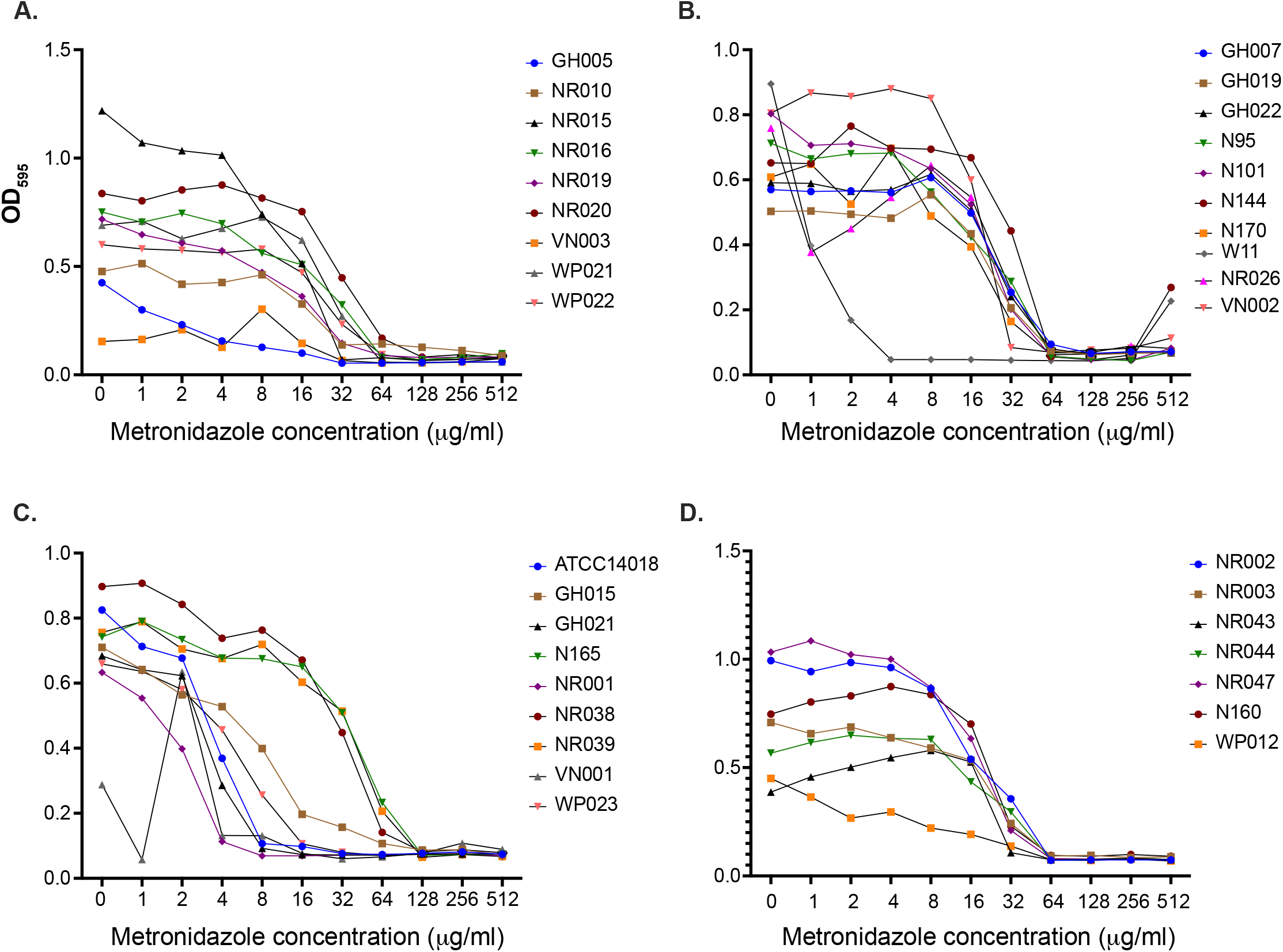
Biofilm formation at different concentrations of metronidazole. Thirty-five *Gardnerella* isolates were grown in media without serum. Each isolate was replicated four times in a 96-well plate. Crystal violet staining was performed to quantify biofilm biomass and OD_595_ measured. Each data point is the average of four technical replicates. Results are shown for isolates in cpn60-defined subgroups A, B, C and D (panels A-D).

## Discussion

It is widely reported that biofilm formation by bacteria provides protection against antibiotics (Costerton 1999; Mah 2012; Lebeaux, Ghigo and Beloin 2014; Bowler, Murphy and Wolcott 2020; Li *et al*. 2020). It has also been suggested that bacterial vaginosis treatment failure and recurrence is largely because of the capacity of *Gardnerella* and other vaginal bacteria to form biofilm (Machado *et al*. 2015; Muzny and Schwebke 2015). Although it has been reported that MIC of metronidazole differs between biofilm and planktonic cultures, the study used only ten isolates of one species of *Gardnerella* (*G. vaginalis*) and did not propose mechanisms of resistance (Li *et al*. 2020).

To promote biofilm or planktonic growth in our study, we used media with or without horse serum. Although the mechanisms are yet to be fully understood, there are reports that inclusion of serum discourages biofilm formation (Abraham and Jefferson 2010; Hammond *et al*. 2010), and that low molecular weight proteins present in the serum may inhibit the transcription of biofilm genes (Abraham and Jefferson 2010). Also, it has been proposed that for motile species, serum may promote twitching motility, which may encourage planktonic growth (Hammond *et al*. 2010). In our study, planktonic growth was dominant in the presence of serum while biofilm growth was dominant in media without serum (Fig. 1). Despite this dramatic difference in growth habit, MIC values in the two conditions were highly correlated and identical MIC values were observed for most isolates regardless of growth medium (Table 1). The MIC of the majority isolates (54.2%) in both media was 64 μg/ml, which is in agreement with a previous study which also reported that the MIC of 50% of tested *Gardnerella* isolates was 64 μg/ml (Petrina *et al*. 2017). The lack of statistically significant difference in MIC between biofilm promoting (media lacking serum) and planktonic growth promoting culture conditions (media containing serum) is perhaps not surprising since in broth microdilution assays, bacteria are exposed to the antibiotic at the time of plate inoculation, before they have the opportunity to form biofilm. These results further suggest that resistance to metronidazole and differences in MIC among isolates are the result of properties of the individual isolates and not solely a function of growth mode (planktonic or biofilm).

Although it has been widely accepted that biofilm formation is a stress response that protects bacteria from insults, including antibacterial compounds, it has also been suggested that biofilm may simply be a default mode of growth for many bacterial species in particular environments (Jefferson 2004). Based on observations of clinical specimens, this certainly seems to be the case for *Gardnerella* in the vaginal microbiome (Swidsinski *et al*. 2005, 2013; Hardy *et al*. 2015, 2016). While the broth microdilution assay and MIC determination provides some information about susceptibility of isolates to metronidazole, it does not simulate the situation *in vivo* where BV associated biofilm is well established prior to treatment. Biofilm formation can protect bacterial cells by the production of extracellular polymeric substances, by release of antibiotic modifying enzymes and extracellular DNA, and reduction of metabolic activity and growth rate (Mah and O’Toole 2001; Hall and Mah 2017). Since metronidazole, an antibiotic bactericidal to a broad range of anaerobes, is a common choice of treatment for BV, we investigated if established *Gardnerella* biofilm can protect its inhabitants from being killed by metronidazole.

To understand how and to what extent biofilm can protect *Gardnerella* cells from metronidazole, 48 h old *Gardnerella* biofilms were exposed to 128 μg/ml metronidazole for 24 h, a concentration higher than the maximum recorded MIC values of isolates we tested. We measured viability and metabolic activity by total viable count and resazurin assay. Resazurin is a high-throughput assay to assess metabolic activity *in vitro* (Osaka and Hefty 2013; Haney *et al*. 2018; Costa *et al*. 2021) and has been used to assess metabolic activity of bacteria residing in biofilms (Mariscal *et al*. 2009; Dalecki, Crawford and Wolschendorf 2016). Our results reveal that metabolic activity is reduced in most of the tested isolates after exposure to metronidazole for 24 h (Fig 3), however, in most cases we tested there was no significant reduction in viable *Gardnerella* in treated biofilms (Fig 2). Our findings suggest that biofilms can protect cells from the killing effect of metronidazole at the cost of reduced metabolic activity. Metronidazole enters bacteria by passive diffusion as a prodrug and has limited activity until it is reduced, which occurs within bacterial cells (Freeman, Klutman and Lamp 1997; Samuelson 1999), and thus, metabolic activity is necessary for the bactericidal effects of the drug to occur. Exposure to antibiotics can also cause the emergence of persister cells in a biofilm, which can survive antibiotic treatment by reducing metabolic activity (Lewis 2008). Reduction of metabolic activity at antibiotic concentrations exceeding the MIC has also been demonstrated in *E. coli* and *S. aureus*; however, in this study metabolic activity was also increased at sub-MIC concentrations (Lobritz *et al*. 2015).

Subinhibitory concentrations of antibiotics can increase the production of extracellular matrix, enhancing biofilm biomass, slowing the diffusion of antibiotics, and reducing exposure of the bacteria within the biofilm (Mah and O’Toole 2001; Mah 2012). We did not observe any enhancement of biofilm formation in the vast majority of the tested *Gardnerella* isolates at sub-MIC concentrations of metronidazole (Fig 4). Enhancement of biofilm formation by sub-inhibitory concentrations of antibiotics likely depends on the mechanisms of actions of antibiotics. Yu et al. demonstrated enhancement of biofilm formation in *Enterococcus faecalis* (a host associated Gram-positive cocci often associated with nosocomial infection) in response to cell wall synthesis inhibitors such as ampicillin, oxacillin, and Fosfomycin, but not in response to protein synthesis, DNA synthesis, and RNA synthesis inhibitors, such as erythromycin, ciprofloxacin, and rifampicin (Yu, Hallinen and Wood 2018). It has been suggested that antimicrobial compounds, such as pyocins, which kill bacteria by damaging the cell wall, can enhance cellular attachment at sub-lethal concentrations leading to increased biofilm formation (Oliveira *et al*. 2015b). Metronidazole, however, kills primarily anaerobic and facultative anaerobic bacteria by formation of reactive oxygen species (ROS), which damages bacterial DNA (Sigeti, Guiney and Davis 1983). Therefore, it is conceivable that due to its mode of action, metronidazole would not be expected to stimulate biofilm formation.

## Conclusions

Metronidazole MICs for individual isolates were found to be unchanged when the isolates were grown in primarily biofilm or primarily planktonic modes. No significant reduction in viable cell count was observed for most *Gardnerella* isolates in established biofilms treated with metronidazole at levels above the MIC, but metabolic activity was reduced in treated biofilms compared to untreated controls. Sub-MIC levels of metronidazole did not stimulate biofilm formation in any isolates. By reducing metabolic activity, *Gardnerella* growing in established biofilms in the vaginal microbiome may avoid the bactericidal effects of metronidazole and resume growth once treatment ends. From a clinical perspective, management of recurrent vaginosis may be improved by methods to mitigate the ecological processes that lead to expansion of *Gardnerella* populations and establishment of biofilm.

## Acknowledgements

This research was supported by a Natural Sciences and Engineering Research Council of Canada Discovery Grant to JEH. The authors are grateful to Champika Fernando for excellent technical support. Thanks to Dr. Joe Rubin, and to the entire Hill Lab for helpful discussions and feedback. No thanks to COVID-19.

## SUPPLEMENTAL MATERIAL

**Fig S1:**
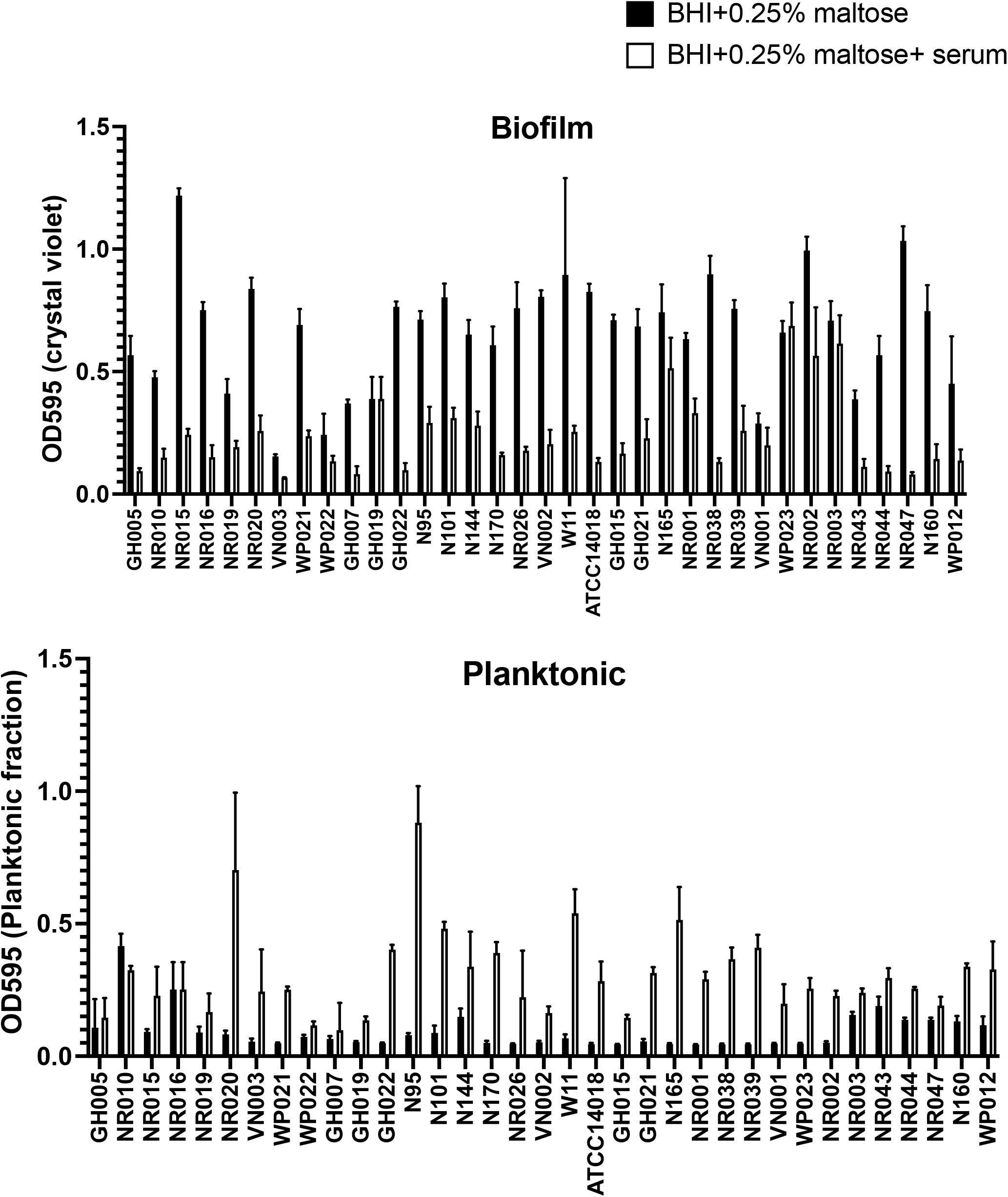
Growth mode of individual isolates is affected by serum. (Upper panel) Biofilm formation in media with serum (BHI+ 0.25% (w/v) maltose + 10% (v/v) heat inactivated horse serum) and without serum (BHI+ 0.25% (w/v) maltose) measured by crystal violet stain. (Lower panel) Planktonic growth in media with serum (BHI+ 0.25% maltose (w/v) + 10% (v/v) heat inactivated horse serum) and without serum (BHI+ 0.25% (w/v) maltose) measured by absorbance at 595 nm. Each bar represents the average of four technical replicates.

## References

Abraham NM, Jefferson KK. A low molecular weight component of serum inhibits biofilm formation in Staphylococcus aureus. Microbial Pathogenesis 2010;49:388–91.

Ahmed A, Earl J, Retchless A et al. Comparative genomic analyses of 17 clinical isolates of Gardnerella vaginalis provide evidence of multiple genetically isolated clades consistent with subspeciation into genovars. J Bacteriol 2012;194:3922–37.

Alauzet C, Lozniewski A, Marchandin H. Metronidazole resistance and nim genes in anaerobes: A review. Anaerobe 2019;55:40–53.

Andersson DI, Hughes D. Microbiological effects of sublethal levels of antibiotics. Nat Rev Microbiol 2014;12:465–78.

Bowler P, Murphy C, Wolcott R. Biofilm exacerbates antibiotic resistance: Is this a current oversight in antimicrobial stewardship? Antimicrobial Resistance & Infection Control 2020;9:162.

Costa P, Gomes ATPC, Braz M et al. Application of the Resazurin Cell Viability Assay to Monitor Escherichia coli and Salmonella Typhimurium Inactivation Mediated by Phages. Antibiotics 2021;10:974.

Costerton JW. Bacterial Biofilms: A Common Cause of Persistent Infections. Science 1999;284:1318–22.

Dalecki AG, Crawford CL, Wolschendorf F. Targeting Biofilm Associated Staphylococcus aureus Using Resazurin Based Drug-susceptibility Assay. JoVE (Journal of Visualized Experiments) 2016:e53925.

Donlan RM, Costerton JW. Biofilms: survival mechanisms of clinically relevant microorganisms. Clin Microbiol Rev 2002;15:167–93.

Freeman CD, Klutman NE, Lamp KC. Metronidazole. Drugs 1997;54:679–708.

Hall CW, Mah T-F. Molecular mechanisms of biofilm-based antibiotic resistance and tolerance in pathogenic bacteria. FEMS Microbiology Reviews 2017;41:276–301.

Hammond A, Dertien J, Colmer-Hamood JA et al. Serum Inhibits P. aeruginosa Biofilm Formation on Plastic Surfaces and Intravenous Catheters. Journal of Surgical Research 2010;159:735–46.

Haney E, Trimble M, Cheng J et al. Critical Assessment of Methods to Quantify Biofilm Growth and Evaluate Antibiofilm Activity of Host Defence Peptides. Biomolecules 2018;8:29.

Hardy L, Cerca N, Jespers V et al. Bacterial biofilms in the vagina. Research in Microbiology 2017;168:865–74.

Hardy L, Jespers V, Abdellati S et al. A fruitful alliance: the synergy between Atopobium vaginae and Gardnerella vaginalis in bacterial vaginosis-associated biofilm. Sexually Transmitted Infections 2016;92:487–91.

Hardy L, Jespers V, Dahchour N et al. Unravelling the bacterial vaginosis-associated biofilm: A multiplex Gardnerella vaginalis and Atopobium vaginae fluorescence in situ hybridization assay using peptide nucleic acid probes. PLoS One 2015;10:e0136658.

Harwich MD, Alves JM, Buck GA et al. Drawing the line between commensal and pathogenic Gardnerella vaginalis through genome analysis and virulence studies. BMC Genomics 2010;11:375.

Hoffman LR, D’Argenio DA, MacCoss MJ et al. Aminoglycoside antibiotics induce bacterial biofilm formation. Nature 2005;436:1171–5.

Jefferson KK. What drives bacteria to produce a biofilm? FEMS Microbiology Letters 2004;236:163–73.

Khan S, Voordouw MJ, Hill JE. Competition among Gardnerella subgroups from the human vaginal microbiome. Front Cell Infect Microbiol 2019;9:374.

Lebeaux D, Ghigo J-M, Beloin C. Biofilm-related infections: bridging the gap between clinical management and fundamental aspects of recalcitrance toward antibiotics. Microbiol Mol Biol Rev 2014;78:510–43.

Lewis K. Multidrug Tolerance of Biofilms and Persister Cells. In: Romeo T (ed.). Bacterial Biofilms. Vol 322. Berlin, Heidelberg: Springer, 2008, 107–31.

Li T, Zhang Z, Wang F et al. Antimicrobial Susceptibility Testing of Metronidazole and Clindamycin against Gardnerella vaginalis in Planktonic and Biofilm Formation. Can J Infect Dis Med Microbiol 2020;2020:1361825.

Lobritz MA, Belenky P, Porter CBM et al. Antibiotic efficacy is linked to bacterial cellular respiration. PNAS 2015;112:8173–80.

Machado D, Castro J, Palmeira-de-Oliveira A et al. Bacterial vaginosis biofilms: Challenges to current therapies and emerging solutions. Frontiers in Microbiology 2015;6:1528.

Mah T-FC. Biofilm-specific antibiotic resistance. Future Microbiology 2012;7:1061–72.

Mah T-FC, O’Toole GA. Mechanisms of biofilm resistance to antimicrobial agents. Trends in Microbiology 2001;9:34–9.

Mariscal A, Lopez-Gigosos RM, Carnero-Varo M et al. Fluorescent assay based on resazurin for detection of activity of disinfectants against bacterial biofilm. Appl Microbiol Biotechnol 2009;82:773–83.

Muzny CA, Schwebke JR. Biofilms: An Underappreciated Mechanism of Treatment Failure and Recurrence in Vaginal Infections. Clinical Infectious Diseases 2015;61:601–6.

Oliveira NM, Martinez-Garcia E, Xavier J et al. Biofilm formation as a response to ecological competition. PLoS Biology 2015a;13:e1002191.

Oliveira NM, Martinez-Garcia E, Xavier J et al. Biofilm Formation As a Response to Ecological Competition. PLOS Biology 2015b;13:e1002191.

Osaka I, Hefty PS. Simple Resazurin-Based Microplate Assay for Measuring Chlamydia Infections. Antimicrobial Agents and Chemotherapy 2013;57:2838–40.

Paramel Jayaprakash T, Schellenberg JJ, Hill JE. Resolution and characterization of distinct cpn60-based subgroups of Gardnerella vaginalis in the vaginal microbiota. PLoS ONE 2012;7:e43009.

Patterson JL, Stull-Lane A, Girerd PH et al. Analysis of adherence, biofilm formation and cytotoxicity suggests a greater virulence potential of Gardnerella vaginalis relative to other bacterial-vaginosis-associated anaerobes. Microbiology 2010;156:392–9.

Petrina MAB, Cosentino LA, Rabe LK et al. Susceptibility of bacterial vaginosis (BV)-associated bacteria to secnidazole compared to metronidazole, tinidazole and clindamycin. Anaerobe 2017;47:115–9.

Raghupathi PK, Liu W, Sabbe K et al. Synergistic Interactions within a Multispecies Biofilm Enhance Individual Species Protection against Grazing by a Pelagic Protozoan. Frontiers in Microbiology 2018;8:2649.

Samuelson J. Why Metronidazole Is Active against both Bacteria and Parasites. Antimicrobial Agents and Chemotherapy 1999;43:1533–41.

Sigeti JS, Guiney DG Jr, Davis CE. Mechanism of action of metronidazole on Bacteroides fragilis. The Journal of Infectious Diseases 1983;148:1083–9.

Smith A. Metronidazole resistance: a hidden epidemic? Br Dent J 2018;224:403–4.

Swidsinski A, Mendling W, Loening-Baucke V et al. Adherent biofilms in bacterial vaginosis. Obstet Gynecol 2005;106:1013–23.

Swidsinski A, Mendling W, Loening-Baucke V et al. An adherent Gardnerella vaginalis biofilm persists on the vaginal epithelium after standard therapy with oral metronidazole. American Journal of Obstetrics and Gynecology 2008;198:97 e1–6.

Swidsinski A, Verstraelen H, Loening-Baucke V et al. Presence of a polymicrobial endometrial biofilm in patients with bacterial vaginosis. PLoS One 2013;8:e53997.

Vaneechoutte M, Guschin A, Van Simaey L et al. Emended description of Gardnerella vaginalis and description of Gardnerella leopoldii sp. nov., Gardnerella piotii sp. nov. and Gardnerella swidsinskii sp. nov., with delineation of 13 genomic species within the genus Gardnerella. International Journal of Systematic and Evolutionary Microbiology 2019;69:679–87.

Verwijs MC, Agaba SK, Darby AC et al. Impact of oral metronidazole treatment on the vaginal microbiota and correlates of treatment failure. American Journal of Obstetrics & Gynecology 2020;222:157.e1-157.e13.

Vidakovic L, Singh PK, Hartmann R et al. Dynamic biofilm architecture confers individual and collective mechanisms of viral protection. Nat Microbiol 2018;3:26–31.

Wiegand I, Hilpert K, Hancock REW. Agar and broth dilution methods to determine the minimal inhibitory concentration (MIC) of antimicrobial substances. Nat Protoc 2008;3:163–75.

Yu W, Hallinen KM, Wood KB. Interplay between Antibiotic Efficacy and Drug-Induced Lysis Underlies Enhanced Biofilm Formation at Subinhibitory Drug Concentrations. Antimicrobial Agents and Chemotherapy 2018;62:e01603–17.

